# Molecular identification of stone loaches of Choman River system, Tigris Basin, based on the Cytochrome *b* gene with an overview of the Cobitoidea Superfamily

**DOI:** 10.1101/210963

**Authors:** Edris Ghaderi, Hamid Farahmand, Barzan Bahrami Kamangar, Mohammad A. Nematollahi

## Abstract

Molecular data and phylogenetic relationships of four Choman loaches species *(Oxynoemacheilus chomanicus, O. zagrosensis, O. kurdistanicus* and *Turcinoemacheilus kosswigi*) recently morphologically described from western Iran were evaluated with 64 species from the Cobitoidea superfamily based on their cytochrome *b* sequences to exhibit the placement of the Choman loaches species within the Cobitoidea superfamily. A comparative analysis of Kimura-2-parameter (K2P) distances was accomplished using sequence divergences of Cobitoidea to calculate intra and interspecific in superfamily, family and genus taxa. The average intraspecific K2P genetic distances of Choman loaches species was 0.005 whereas this value was 0.016 for the Cobitoidea superfamily. Molecular phylogenetic relationships were assessed using Maximum likelihood and Bayesian methods. Dendrograms obtained by these methods revealed all four Choman loaches species as distinct species among other reported Nemacheilidae Spp. These species were clustered with *Oxynoemacheilus* and *Turcinoemacheilus* genera within other species in the Nemacheilidae family. The phylogenetic analysis revealed that Cobitoidea superfamily consists of nine families ((Gyrinocheilidae + Botiidae) + ((Catostomidae + Vaillentellidae) + ((Nemacheilidae + Cobitidae) + ((Ellopostomidae + Gastromyzontidae) + Balitoridae)))) and indicated Nemacheilidae is a valid and distinct family from Balitoridae.

## Introduction

Loaches of the family Nemacheilidae represent a major lineage of the superfamily Cobitoidea that are diverse in freshwaters of Asia, Europe and north-east Africa with more than 45 genera and 570 known species (Kottelat, 2012). Eight genera and more than 40 species of nemacheilid are widely distributed in Iran’s drainage basins (Coad, 1998; Nalbant & Bianco, 1998; Golzarianpour et al., 2009, 2011 and 2013; Abdoli et al., 2011; Kamangar et al., 2014; Freyhof et al., 2016). Although most of the new species are described morphologically, these approaches are faced with challenges (Robertson et al., 2001). Molecular procedures have been applied in many studies for new species identification and it is clear that these methods have great potential in resolving identification and phylogenic relationship of cryptic species and could be highly effective in classification of confusing species (Teletchea, 2009). Cytochrome *b* gene (Cytb) is frequently utilized for fish identification and phylogenetic study (Song et al., 1998; Biswas et al., 2001; Farias et al., 2001; Perkins & Schall, 2002; Perdices et al., 2004; Baharum & Nurdalila, 2011).

Choman watershed is a sub-basin of Tigris and located in the west of Iran. Recently, four stone loach species from Nemacheilidae family; *Turcinoemacheilus kosswigi* (Banarescu & Nalbant, 1964) and three new species from *Oxynoemacheilus* (*O. chomanicus, O. zagrosensis* and *O. kurdistanicus*), have been reported from this watershed based on morphological and some osteological characters (Kamangar et al., 2014). However, molecular data on these species have not as yet been reported. In order to bring molecular data to complete introducing and demonstrating the taxonomic status of these loach species from the Choman River sub-basin, molecular tools based on Cytochrome *b* gene was used complementary to morphological approaches in this study.

The taxonomy of Cobitoidea superfamily has been a controversial subject for ichthyologists in the last century. For instance, Nemacheilid loaches were initially classified in the Cobitidae family (Regan, 1911; Hora, 1932) and thereafter transferred to the Balitoridae family (Sawada, 1982; Siebert, 1987). For the first time, Nalbant (2002) changed this classification by treating the Nemacheilinae as the Nemacheilidae family. The results of the first molecular phylogenetic study of Cobitoidea by Liu et al. (2002) in which mtDNA control region sequences was used suggested once again the Nemacheilinae rank for this group within the Cobitoidea. Nevertheless, this classification was not approved by ichthyologists due to inadequate testing of the phylogenetic relationship of loaches sensu stricto (Tang et al., 2006). In following molecular studies, the Nalbant (2002) classification was confirmed and nemacheilid loach fishes were classified in a separate family from the Balitoridae as the Nemacheilidae family (Šlechtová et al., 2007; Bohlen & Šlechtová, 2009; Chen et al., 2009). Finally, Kottelat (2012) identified 10 families (Gyrinocheilidae, Botiidae, Vaillantellidae, Cobitidae, Ellopostomatidae, Barbuccidae, Balitoridae, Gastromyzontidae, Serpenticobitidae and Nemacheilidae) in the Cobitoidea superfamily and Nemacheilidae was introduced as a valid family in this group. However, in some systematic and taxonomic reviews, the “Nemacheilinae” is still used (Wang et al., 2012; Du et al., 2012; Tang et al., 2013; Prokofiev & Golubstove, 2013; Li et al., 2015; Que et al., 2016) and using different expressions for this group has led to confusion.

The objectives of this study were firstly to evaluate cytochrome *b* sequence variation and the morphological tools associated by morphometric truss network analysis in grouping and identifying Choman loach species. Secondly, the aim was to examine the phylogenetic relationships in the superfamily Cobitoidea based on Cytochrome *b* sequences using new taxa and to note the placement of Choman loach species.

## Materials and methods

### Fish sampling

One hundred twenty five loach specimens from the Choman sub-basin including *O. chomanicus* (38.1- 56.2 mm SL, n=40), *O. zagrosensis* (48.4- 58.4 mm SL, n=20), *O. kurdistanicus* (32.2- 60.5 mm SL, n=35) and *T. kosswigi* (25- 49.4 mm SL, n=30) were sampled from the same four locations as reported by Kamangar et al. (2014) between August and September 2011 (Fig. 1). Five specimens of muscle tissue were taken from each species for molecular study.

**Fig. 1.**
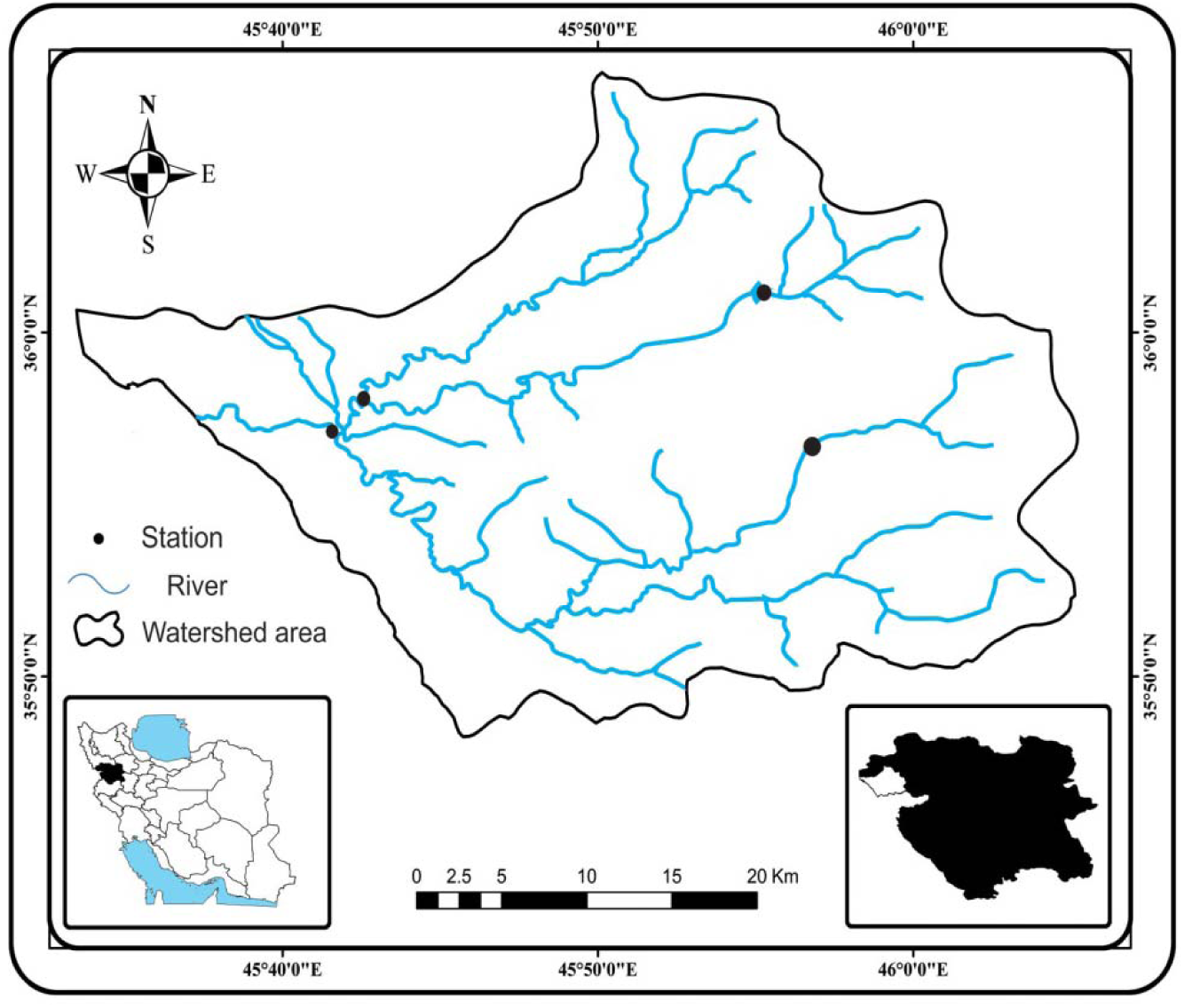
Map of Choman river basin showing the Choman loaches sampling sites.

### Morphological analysis

Truss network is a quantitative method representing the complete shape of fish and is an effective method for obtaining information regarding the great similarity of morphological variation (Cavalcanti et al., 2011). In the truss network system, 16 homologous landmarks (Fig.2) delineating 38 distance characters (Table 1) were measured on the body with calipers following methods outlined by Strauss and Bookstein (1982). An allometric method (Elliott et al., 1995) was used to remove size-dependent variation in morphometric characters: *M_adj_* = M (L_s_/ L_0_) ^b^

**Fig. 2.**
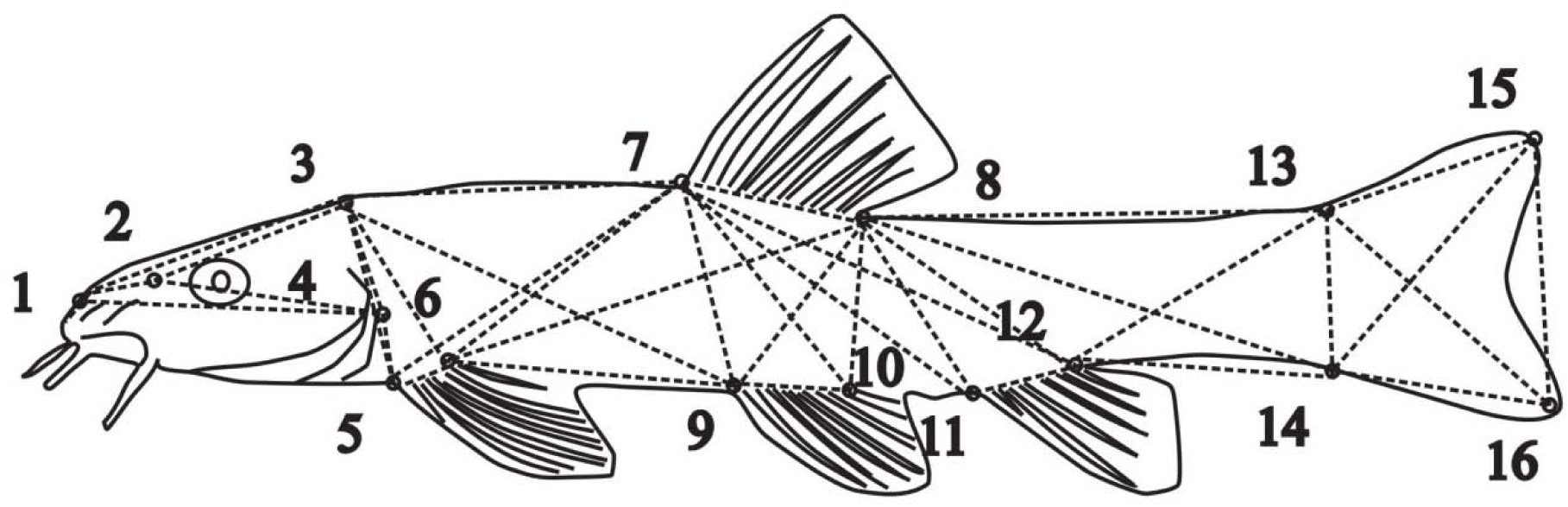
The truss network, anatomical landmark points (16 points) and distance measured for Choman loaches species. Anatomical landmark points, 1: Tip of snout; 2: End of nostril; 3: Forehead (end of frontal bone); 4: Hind most point of operculum; 5: Insertion of pectoral fin; 6: Insertion of last pectoral fin ray; 7: Insertion of dorsal fin; 8: Insertion of last dorsal fin ray; 9: Insertion of pelvic fin; 10: Insertion of last pelvic fin ray; 11: Insertion of anal fin; 12: Insertion of last anal fin ray; 13: Insertion of first caudal fin ray in upper lobe; 14: Insertion of first caudal fin ray in lower lobe; 15: The most distal point of caudal fin ray in upper lobe; 16:The most distal point of caudal fin ray in lower lobe.

**Table 1.** Comparisons of the landmark distance measurements between Choman loaches species. Values are mean ± standard error. The values that followed by different letters indicate a significant difference between species for the same measurements (P < 0.05).

| Distance measured | Characters abbreviation | *O. chomanicus* | *O. zagrosensis* | *O. kurdistanicus* | *T. kosswigi* |
| --- | --- | --- | --- | --- | --- |
| 1-2 | SPN | 2.844 ± 0.347<sup>b</sup> | 3.519 ± 0.276<sup>c</sup> | 3.034 ± 0.516<sup>b</sup> | 2.308 ± 0.240<sup>a</sup> |
| 1-3 | SOC | 8.601 ± 0.618<sup>b</sup> | 10.084 ± 0.597<sup>c</sup> | 8.863 ± 0.547<sup>b</sup> | 6.517 ± 0.468<sup>a</sup> |
| 1-4 | SPG | 10.374 ± 0.509<sup>c</sup> | 11.746 ± 0.361<sup>d</sup> | 9.815 ± 0.565<sup>b</sup> | 7.241 ± 0.541<sup>a</sup> |
| 2-3 | PNOC | 6.162 ± 0.392<sup>b</sup> | 6.954 ± 0.394<sup>c</sup> | 6.336 ± 0.379<sup>b</sup> | 4.862 ± 0.320<sup>a</sup> |
| 2-4 | PNPG | 8.170 ± 0.589<sup>c</sup> | 9.048 ± 0.469<sup>d</sup> | 7.537 ± 0.329<sup>b</sup> | 5.460 ± 0.314<sup>a</sup> |
| 3-4 | OCPG | 5.945 ± 0.590<sup>c</sup> | 6.725 ± 0.604<sup>d</sup> | 4.923 ± 0.520<sup>b</sup> | 3.178 ± 0.383<sup>a</sup> |
| 3-5 | OCAP | 6.418 ± 0.556<sup>c</sup> | 7.246 ± 0.375<sup>d</sup> | 5.742 ± 0.484<sup>b</sup> | 3.842 ± 0.469<sup>a</sup> |
| 3-6 | OCPD | 7.324 ± 0.557<sup>c</sup> | 8.100 ± 0.326<sup>d</sup> | 6.388 ± 0.625<sup>b</sup> | 4.368 ± 0.568<sup>a</sup> |
| 3-7 | OCAD | 15.042 ± 1.080<sup>b</sup> | 17.298 ± 1.177<sup>c</sup> | 14.251 ± 1.926<sup>a</sup> | 16.791 ± 1.385<sup>c</sup> |
| 3-9 | OCAV | 18.730 ± 0.944<sup>c</sup> | 21.488 ± 0.756<sup>d</sup> | 17.293 ± 1.076<sup>b</sup> | 15.733 ± 0.624<sup>a</sup> |
| 4-5 | PGAP | 1.528 ± 0.394<sup>b</sup> | 1.479 ± 0.478<sup>b</sup> | 1.463 ± 0.395<sup>b</sup> | 1.091 ± 0.352<sup>a</sup> |
| 5-6 | APPP | 2.243 ± 0.337<sup>b</sup> | 2.689 ± 0.301<sup>c</sup> | 2.368 ± 0.456<sup>b</sup> | 1.524 ± 0.255<sup>a</sup> |
| 5-7 | APAD | 13.767 ± 1.370<sup>a</sup> | 16.149 ± 0.830<sup>c</sup> | 13.647 ± 1.628<sup>a</sup> | 15.197 ± 0.997<sup>b</sup> |
| 6-7 | PPAD | 13.106 ± 1.378<sup>a</sup> | 14.910 ± 1.071<sup>b</sup> | 12.820 ± 1.540<sup>a</sup> | 14.786 ± 0.896<sup>b</sup> |
| 6-8 | PPPD | 18.599 ± 1.494<sup>a</sup> | 21.816 ± 0.802<sup>b</sup> | 18.365 ± 2.002<sup>a</sup> | 18.677 ± 0.694<sup>a</sup> |
| 6-9 | PPAV | 13.224 ± 1.216<sup>b</sup> | 14.998 ± 0.969<sup>c</sup> | 12.288 ± 1.080<sup>a</sup> | 13.075 ± 2.139<sup>b</sup> |
| 7-8 | ADPD | 6.473 ± 0.551<sup>b</sup> | 8.349 ± 0.672<sup>d</sup> | 6.837 ± 0.776<sup>c</sup> | 4.148 ± 0.513<sup>a</sup> |
| 7-9 | ADAV | 7.373 ± 0.840<sup>c</sup> | 9.012 ± 0.955<sup>d</sup> | 6.805 ± 0.652<sup>b</sup> | 4.651 ± 0.516<sup>a</sup> |
| 7-10 | ADPV | 7.605 ± 1.061<sup>c</sup> | 8.506 ± 0.812<sup>d</sup> | 6.745 ± 0.786<sup>b</sup> | 4.658 ± 0.434<sup>a</sup> |
| 7-11 | ADAA | 15.353 ± 0.844<sup>c</sup> | 17.808 ± 0.982<sup>d</sup> | 13.336 ± 0.964<sup>b</sup> | 10.432 ± 1.071<sup>a</sup> |
| 7-12 | ADPA | 18.115 ± 0.834<sup>c</sup> | 21.346 ± 1.068<sup>d</sup> | 16.088 ± 0.933<sup>b</sup> | 12.969 ± 0.668<sup>a</sup> |
| 8-9 | PDAV | 7.891 ± 0.759<sup>b</sup> | 9.837 ± 1.011<sup>c</sup> | 7.725 ± 1.101<sup>b</sup> | 6.857 ± 0.733<sup>a</sup> |
| 8-10 | PDPV | 7.781 ± 1.030<sup>b</sup> | 9.354 ± 0.632<sup>c</sup> | 7.387 ± 1.369<sup>b</sup> | 6.658 ± 0.886<sup>a</sup> |
| 8-11 | PDAA | 9.094 ± 0.772<sup>c</sup> | 10.618 ± 0.402<sup>d</sup> | 6.973 ± 0.465<sup>b</sup> | 6.418 ± 0.739<sup>a</sup> |
| 8-12 | PDPA | 11.347 ± 0.751<sup>c</sup> | 13.143 ± 0.828<sup>d</sup> | 9.116 ± 0.705<sup>b</sup> | 8.378 ± 0.633<sup>a</sup> |
| 8-13 | PDFLL | 16.639 ± 0.821<sup>b</sup> | 19.068 ± 0.924<sup>c</sup> | 16.260 ± 0.893<sup>b</sup> | 14.436 ± 0.725<sup>a</sup> |
| 8-14 | PDFUL | 17.670 ± 0.850<sup>c</sup> | 20.290 ± 0.876<sup>d</sup> | 16.682 ± 0.775<sup>b</sup> | 14.879 ± 0.615<sup>a</sup> |
| 9-10 | AVPV | 1.934 ± 0.216<sup>b</sup> | 2.290 ± 0.182<sup>c</sup> | 1.989 ± 0.281<sup>b</sup> | 1.452 ± 0.352<sup>a</sup> |
| 10-11 | PVAA | 9.462 ± 0.580<sup>b</sup> | 10.837 ± 0.696<sup>c</sup> | 7.942 ± 0.721<sup>a</sup> | 9.429 ± 1.353<sup>b</sup> |
| 11-12 | AAPA | 3.935 ± 0.505<sup>b</sup> | 4.494 ± 0.428<sup>c</sup> | 4.105 ± 0.509<sup>b</sup> | 3.152 ± 0.410<sup>a</sup> |
| 12-13 | PAFLL | 8.758 ± 0.607<sup>b</sup> | 10.584 ± 0.509<sup>d</sup> | 9.790 ± 0.597<sup>c</sup> | 7.508 ± 0.448<sup>a</sup> |
| 12-14 | PAFUL | 7.334 ± 0.687<sup>b</sup> | 8.429 ± 0.792<sup>c</sup> | 9.355 ± 0.695<sup>d</sup> | 6.944 ± 0.457<sup>a</sup> |
| 13-14 | FLLFUL | 4.966 ± 0.589<sup>c</sup> | 6.172 ± 0.487<sup>d</sup> | 3.997 ± 0.288<sup>b</sup> | 3.195 ± 0.237<sup>a</sup> |
| 13-15 | FLLEUL | 8.970 ± 0.670<sup>b</sup> | 10.161 ± 0.653<sup>c</sup> | 10.217 ± 0.565<sup>c</sup> | 6.321 ± 0.432<sup>a</sup> |
| 13-16 | FLLELL | 8.585 ± 0.462<sup>b</sup> | 9.583 ± 0.423<sup>c</sup> | 9.570 ± 0.787<sup>c</sup> | 6.121 ± 0.422<sup>a</sup> |
| 14-15 | FULEUL | 8.504 ± 0.447<sup>b</sup> | 9.348 ± 0.515<sup>c</sup> | 9.698 ± 0.736<sup>d</sup> | 6.237 ± 0.448<sup>a</sup> |
| 14-16 | FULELL | 9.752 ± 0.742<sup>b</sup> | 11.071 ± 0.722<sup>d</sup> | 10.230 ± 0.849<sup>c</sup> | 6.564 ± 0.425<sup>a</sup> |
| 15-16 | EULELL | 4.158 ± 0.860<sup>c</sup> | 4.609 ± 1.121<sup>c</sup> | 3.606 ± 1.075<sup>b</sup> | 2.853 ± 0.520<sup>a</sup> |

In this equation *M* is the original measurement, *M_adj_* is the size adjusted measurement, *L_0_* is the standard length of the fish, *L_s_* is the overall mean of the standard length for all fish from all samples in each analysis. The coefficient of *b* was estimated for each character from the observed data as the slope of the regression of *log M* on *log L_0_* using all fish in the groups. Analysis of variance (ANOVA) followed by Duncan’s multiple range test was applied to test any significant differences of landmarks between the four Choman loach species. The size-adjusted data were submitted to a stepwise discriminant function analysis (DFA). Squared Mahalanobis distances matrix was obtained from the cluster analysis and used to draw a dendrogram based on the unweighted pair group method analysis (UPGMA) (Bektaz & Belduz, 2009). Statistical analysis was performed using SPSS ver. 16 and STATISTICA ver. 6.0 software.

### DNA extraction and Cytochrome *b* sequencing

DNA was extracted using salting out protocol from muscle tissue (Cawthorn et al., 2011). To amplify a 1140 bp of Cytochrome *b*, DNA extracts were subjected to PCR amplification, using primer set of L14724 (5’-GAC TTG AAA AAC CAC CGT TG-3’) and H15915 (5’-CTC CGA TCT CCG GAT TAC AAG AC-3’) (Tang *et al.,* 2006). PCR was performed at an initial denaturation step at 94 °C for 3 min followed by 35 cycles at 94 °C for 30 s, 52-58 °C for 45 s, 72 °C for 1 min and a final extension at 72 °C for 8 min. The amplified fragments were purified with Promega DNA purification kit following the manufacturer’s instructions. The purified fragments were sequenced bidirectional by BIONEER Company. All sequences are available from the GenBank database.

### Phylogenetic analysis

For phylogenetic analysis, sequences from Choman loaches species (20 sequences) were evaluated with sequences from 180 individuals representing a total of 64 species from Cobitoidea. These sequences were downloaded from GenBank. Cytochrome *b* sequence of *Capoeta trutta* (JF798333) and *Cyprinus carpio* (AB158807) were used as out-grouping in this study. We followed the classification of Cobitoidea proposed by Kottelat (2012). Moreover, we evaluated the Catostomidae relationship with Cobitoidea as proposed by Šlechtová et al. (2007) as a family member of Cobitoidea. There was no reported Cytochrome *b* sequences from Barbuccidae and Serpenticobitidae families and thus sequences from Gyrinocheilidae, Botiidae, Vaillentellidae, Nemacheilidae, Cobitidae, Gastromyzontidae, Catostomidae, Ellopostomidae and Balitoridae families only were used in this analysis.

Kimura two parameter (K2P) distance model (Kimura, 1980) was used to calculate sequence divergences in the Cobitoidea superfamily. Several categories of K2P were calculated to test the first objective of this study: intra and interspecific distances in the Cobitoidea superfamily, intra and interspecific distances in the Balitoridae, Nemacheilidae, Catostomidae, Cobitidae, Botiidae and Gastromyzontidae family, intra and interspecific distances for *Oxynoemacheilus* genus and finally intraspecific distance in Choman loach species. The averages of each category were compared statistically to test any possible overlapping between these distances. K2P genetic distances were calculated using MEGA 4.0 software package (Tamura et al., 2007). The comparisons between the average genetic distances were carried out in XLSTAT software trial version. The Mann-Whitney test was used to statistically compare all intraspecific and interspecific comparisons. To calculate conspecific (intraspecific) distances, the species used were ones that were represented at least by two or more specimens in the data bank. Confamilial (intraspecific distance in each family) distances were also calculated when at least two genera sequences were available.

Phylogenetic trees were reconstructed for the gene data set using Maximum Likelihood (ML) and Bayesian Analyses (BA). Sequences were aligned using Clustal X ver. 1.85 (Thompson et al., 1997). Phylogenetic trees using ML were estimated as implemented in RAxML (Randomized Axelerated Maximum Likelihood, version 7.0.4) (Stamatakis, 2006). The Akaike information criterion (AIC) implemented in MODELTEST v3.4 (Posada and Crandall, 1998) was used to identify the optimal molecular evolutionary model for each partition on the sequence data set. GTR+I+G model were utilized to obtain the optimal ML trees and bootstrap support. Robustness of the inferred trees was evaluated using bootstrap analysis on 1,000 pseudoreplications using RAxML 7.0.4 (Felsenstein, 1985; Stamatakis et al., 2008). BA analyses were conducted using Mr. Bayes v3.1.2 (Huelsenbeck and Ronquist, 2001). For BA, 5,000,000 cycles were implemented for four simultaneous Monte Carlo Markov chains, sampling the Markov chain at intervals of 100 generations. Log-likelihood stability was attained after 100,000 generations; the first 1,000 trees were discarded as “burn-in” in each analysis. Support for BI tree nodes was determined based on values of Bayesian posterior probabilities. Final trees were visualized in the FigTree v.1.4.2 program (Rambaut, 2009).

## Results

### Morphological traits

The Choman loach species were well distinguished morphometrically based on truss network analysis. The ANOVA results showed significant differences between species for all truss distances (p < 0.01). The effect of sex was not significant within the same species for all examined measurements. The comparison of mean truss distances between the four loach species revealed significant differences between them (Table 1). Most landmarks were significantly longer in *O. zagrosensis* than other Choman loach species (Table 1).

Fourteen landmarks (ADPA, PAFUL, OCPP, OCAD, SPG, FULEUL, PVAA, OCPG, PPAV, PDPA, ADPD, EULELL, OCAV, and FLLELL) were revealed by stepwise DFAs as the most discriminative landmarks to differentiate the loach species. The canonical discriminant function analysis produced three functions; the first two canonical discriminant functions explained 95% of variability between species (Table 2). The Wilks’ lambda statistics showed the power of discriminant functions in separating species (Table 3). The group centroid of each species was comprehensively disparate based on visual analysis of scattering diagrams (Fig. 3).

**Table 2.** Eigenvalue, percentage of variance and cumulative variance explained by three discriminant functions in the truss network analysis of Choman loaches species.

| Function | Eigenvalue | % of Variance | Cumulative % |
| --- | --- | --- | --- |
| 1 | 36.198 | 75.5 | 75.5 |
| 2 | 9.345 | 19.5 | 95.0 |
| 3 | 2.420 | 5.0 | 100.0 |

**Table 3.** Test of significance of discriminant functions for verifying differences of Choman loach species.

| Functions | Wilks' Lambda | Chi-square | df | P value |
| --- | --- | --- | --- | --- |
| 1 through 3 | 0.001 | 825.962 | 42 | 0.000 |
| 2 through 3 | 0.028 | 410.093 | 26 | 0.000 |
| 3 | 0.292 | 141.393 | 12 | 0.000 |

**Fig. 3.**
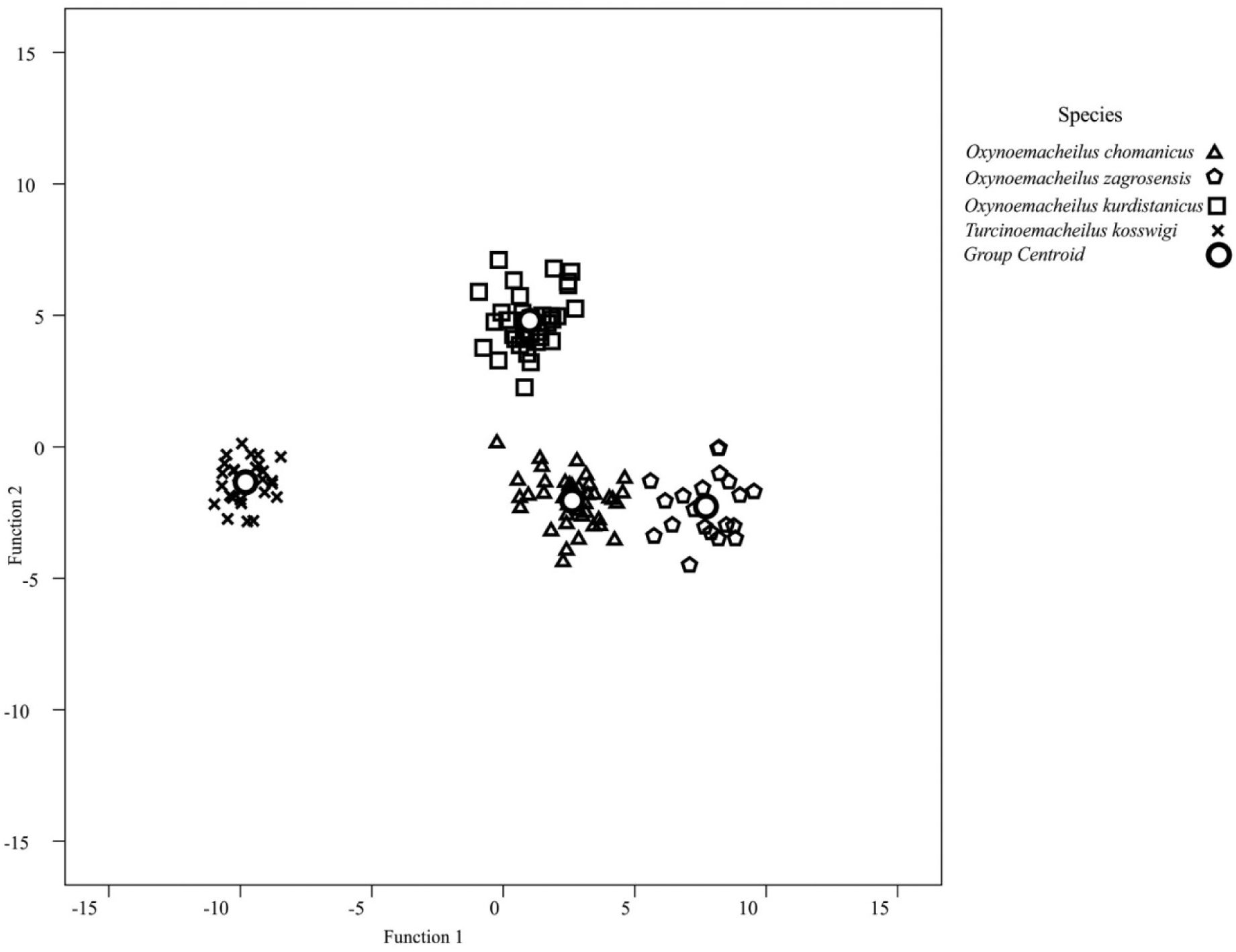
Scatterplot of individual distances measured based on truss network analysis from the first two canonical functions for Choman loaches species.

Cross validation results showed that 99.2% of individuals were correctly classified in their species groups based on morphological differences (Table 4) and only one specimen from *O. zagrosensis* was incorrectly classified in the *O. chomanicus* group (Fig. 3). The dendrogram derived from cluster analysis of groups of centroids showed two separate groups: one included the *Oxynoemacheilus spp.* and the other included the *Turcinoemacheilus* (Fig. 4). These groupings confirm the results obtained from the discriminant analysis for these species.

**Table 4.**
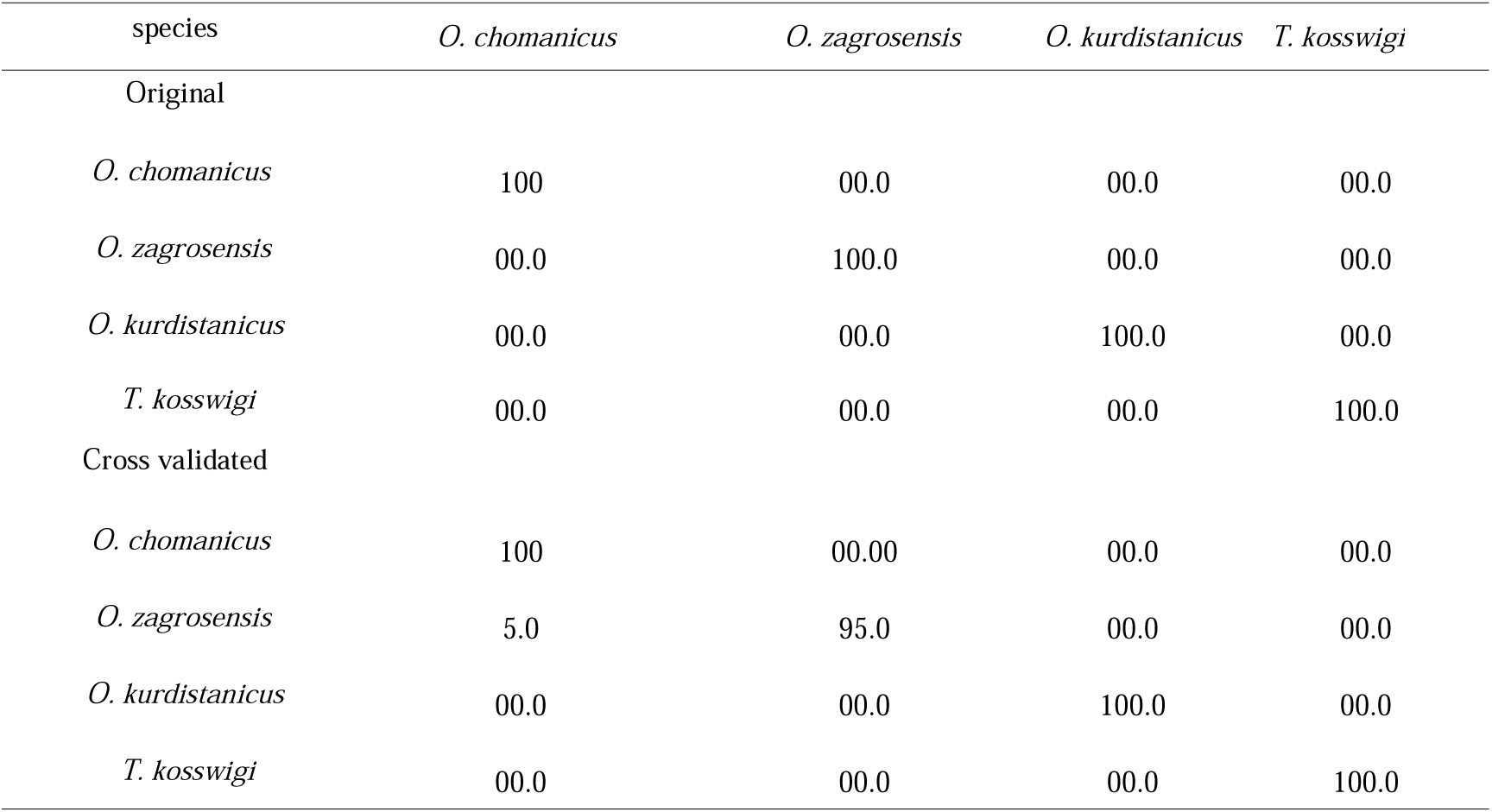
Percentage of specimens classified in each species and after cross validation for truss network distance measurements for Choman loaches species.

| species | *O. chomanicus* | *O. zagrosensis* | *O. kurdistanicus* | *T. kosswigi* |
| --- | --- | --- | --- | --- |
| Original |  |  |  |  |
| *O. chomanicus* | 100 | 00.0 | 00.0 | 00.0 |
| *O. zagrosensis* | 00.0 | 100.0 | 00.0 | 00.0 |
| *O. kurdistanicus* | 00.0 | 00.0 | 100.0 | 00.0 |
| *T. kosswigi* | 00.0 | 00.0 | 00.0 | 100.0 |
| Cross validated |  |  |  |  |
| *O. chomanicus* | 100 | 00.00 | 00.0 | 00.0 |
| *O. zagrosensis* | 5.0 | 95.0 | 00.0 | 00.0 |
| *O. kurdistanicus* | 00.0 | 00.0 | 100.0 | 00.0 |
| *T. kosswigi* | 00.0 | 00.0 | 00.0 | 100.0 |

**Fig. 4.**
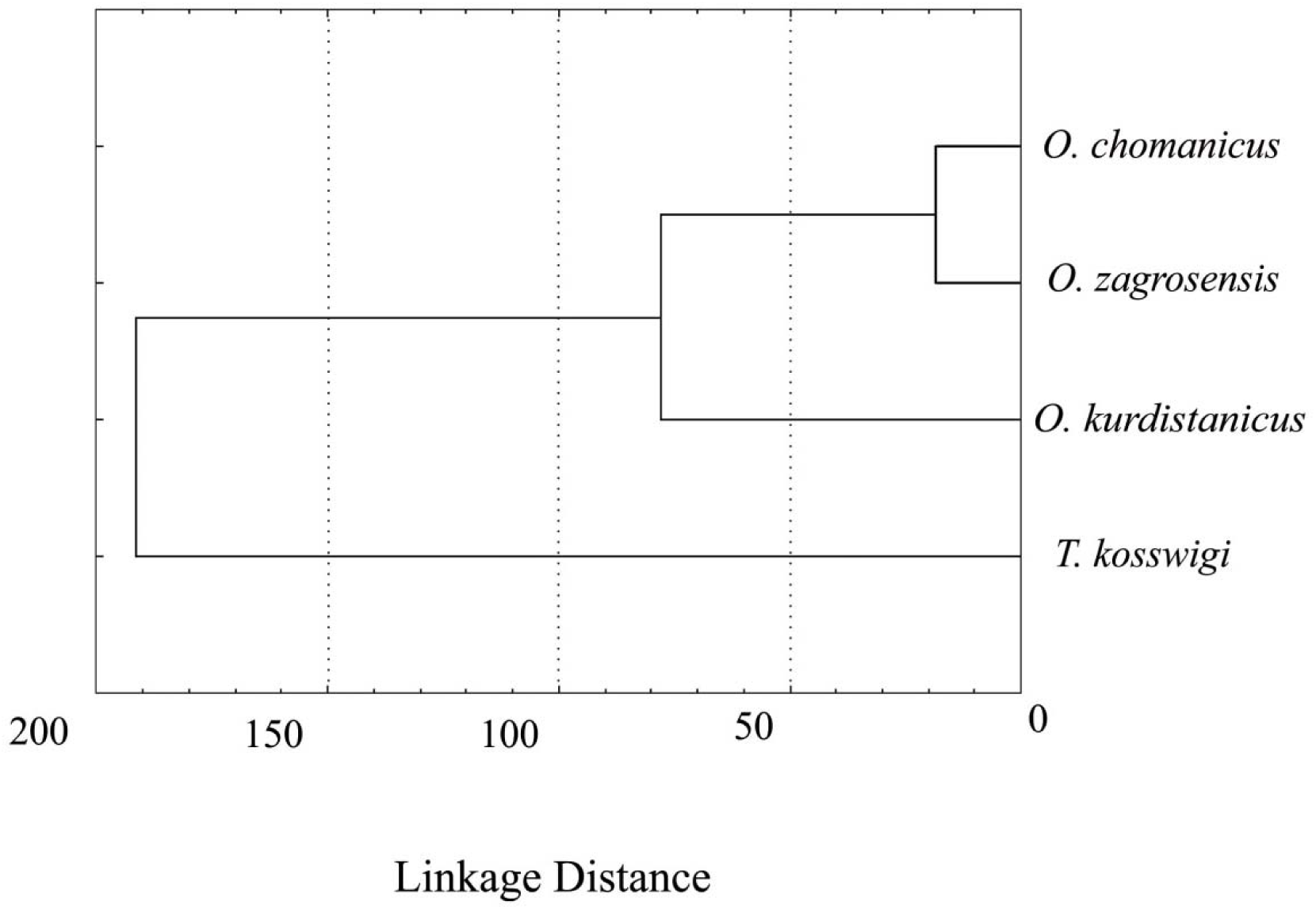
Dendrogram derived from cluster analysis of Mahalanobis distance using UPGMA between Choman loaches species.

### Molecular study

The average sequence length was 1097 bp (range= 1011 to 1131 bp); all read lengths were greater than 1000 bp. No internal indels were observed in any sequences generated herein or downloaded from the Genbank. There were 1157 nucleotide sites, of which 685 were parsimony informative sites (55%). The average nucleotide composition of *O. chomanicus, O. zagrosensis, O. kurdistanicus* and *T. kosswigi* sequences were Adenine (A) 25%, 24.9%, 25.4% and 28.7%, Thymine (T) 31%, 30.4%, 30.2% and 31.5%, Cytosine (C) 27.5%, 27.2%, 27.8% and 24.4% and Guanine (G) 16.5%, 17.4%, 16.6% and 15.4%, respectively. In all four species, the content of A+T was higher than that of C+G. Transitions (Ti) / transversions (Tv) ratios for *O. chomanicus, O. zagrosensis, O. kurdistanicus* and *T. kosswigi* were 2.54, 1.28, 0.79 and 2.19, respectively. The average nucleotide composition in Cobitoidea superfamily was Adenine (A) 26.3%, Thymine (T) 30.7%, Cytosine (C) 27.2% and Guanine (G) 15.8%, and Transitions (Ti) / transversions (Tv) ratios in this superfamily was estimated as 1.82.

The K2P distances were calculated in the Cobitoidea superfamily and Nemacheilidae, Balitoridae, Gastromyzontidae, Cobitidae, Botiidae, Catostomidae families and the *Oxynoemacheilus* genus in order to reveal any possible overlapping and statistically significant differences between these distances (Table 5). Significant differences were observed between intraspecific and interspecific K2P distances in all comparisons (p < 0.001). There was no consequential overlap between all intra and interspecific K2P distance comparisons (Table 5). The mean interspecific K2P distances were higher by at least 10-fold than their intraspecific genetic distances except those for Balitoridae, Gastromyzontidae and Catostomidae families (Table 5). The average interspecies of *Oxynoemacheilus* genus were 32.50-fold higher than that amongst individuals within (intraspecific) the *Oxynoemacheilus* species (Table 5). This ratio was 32.50, 21.67 and 43.33 higher for *O. chomanicus, O. zagrosensis* and *O. kurdistanicus*, respectively. The average interspecific from different species but within the Nemacheilidae family (0.216) was 54.00 fold higher than that amongst individuals within the *Oxynoemacheilus* species (0.004) (Table 5).

**Table 5.** Summary of intra and interspecific K2P distances in the Cobitoidea superfamily, 6 families, *Oxynoemacheilus* genus and Choman loaches species

| Taxa | Comparisons | Number of comparisons | Median | Minimum | Maximum | Mean | ratio | Standard deviation | Z, P-value |
| --- | --- | --- | --- | --- | --- | --- | --- | --- | --- |
| Cobitoidea | intraspecific | 306 | 0.006 | 0.000 | 0.108 | 0.016 |  | 0.024 |  |
|  | interspecific | 19664 | 0.239 | 0.002 | 0.339 | 0.231 | 14.44 | 0.044 | -25.96, 0.000 |
| Nemacheilidae | intraspecific | 131 | 0.006 | 0.000 | 0.108 | 0.018 |  | 0.028 |  |
|  | interspecific | 4240 | 0.223 | 0.002 | 0.317 | 0.216 | 12.00 | 0.049 | -19.28, 0.000 |
| Balitoridae | intraspecific | 35 | 0.006 | 0.000 | 0.028 | 0.010 |  | 0.009 |  |
|  | interspecific | 265 | 0.091 | 0.016 | 0.137 | 0.091 | 9.10 | 0.025 | -9.56, 0.000 |
| Gastromyzontidae | intraspecific | 10 | 0.015 | 0.005 | 0.082 | 0.032 |  | 0.034 |  |
|  | interspecific | 26 | 0.157 | 0.041 | 0.182 | 0.119 | 3.72 | 0.058 | -3.64, 0.000 |
| Cobitidae | intraspecific | 45 | 0.005 | 0.000 | 0.075 | 0.012 |  | 0.018 |  |
|  | interspecific | 889 | 0.151 | 0.050 | 0.218 | 0.150 | 12.50 | 0.032 | -11.30, 0.000 |
| Botiidae | intraspecific | 9 | 0.005 | 0.000 | 0.030 | 0.007 |  | 0.009 |  |
|  | interspecific | 57 | 0.153 | 0.073 | 0.190 | 0.144 | 20.57 | 0.034 | -4.79, 0.000 |
| Catostomidae | intraspecific | 6 | 0.004 | 0.000 | 0.090 | 0.021 |  | 0.035 |  |
|  | interspecific | 60 | 0.184 | 0.093 | 0.237 | 0.168 | 8.00 | 0.041 | -4.02, 0.000 |
| Oxynoemacheilus | intraspecific | 43 | 0.003 | 0.000 | 0.010 | 0.004 |  | 0.004 |  |
|  | interspecific | 257 | 0.164 | 0.011 | 0.201 | 0.130 | 32.50 | 0.063 | -10.50, 0.000 |
| *O. chomanicus* | intraspecific | 10 | 0.005 | 0.000 | 0.008 | 0.004 | 32.50 | 0.003 | -5.370, 0.000 |
| *O. zagrosensis* | intraspecific | 10 | 0.006 | 0.000 | 0.008 | 0.006 | 21.67 | 0.003 | -5.370, 0.000 |
| *O. kurdistanicus* | intraspecific | 10 | 0.003 | 0.000 | 0.006 | 0.003 | 43.33 | 0.002 | -5.370, 0.000 |
| *T. kosswigi* | intraspecific | 10 | 0.0097 | 0.006 | 0.016 | 0.009 |  | 0.003 |  |

The same topologies were achieved when the Cobitoidea superfamily data was analyzed with BA (Fig. 5) and ML (Fig. 6) methods. In both analyses, the Choman loach species were correctly classified within their family and genus clades. *O. chomanicus* and *O. zagrosensis* formed monophyletic groups and then gathered in one clade with *O. kurdistanicus.* The recently described Choman loach species were distinguished from other *Oxynoemacheilus* spp. which had their Cytochrome *b* gene sequences reported.

**Fig. 5.**
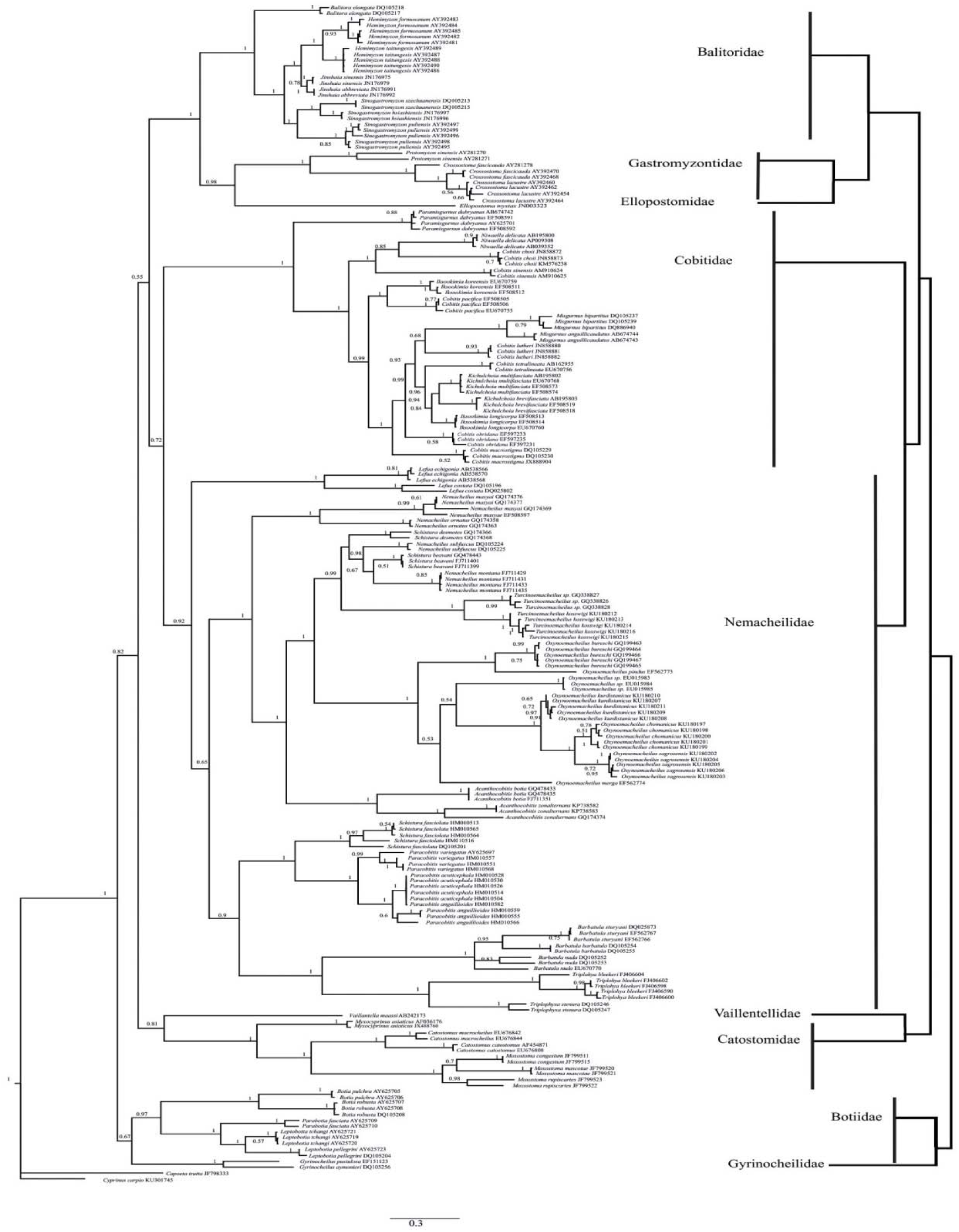
The 50% majority-rule consensus tree resulting from Bayesian analysis of the concatenated cytochrome *b* sequences dataset. Numbers at nodes represent posterior probabilities for Bayesian analysis. An overview of the relationship obtained for Cobitoidea superfamily shown (right).

**Fig. 6.**
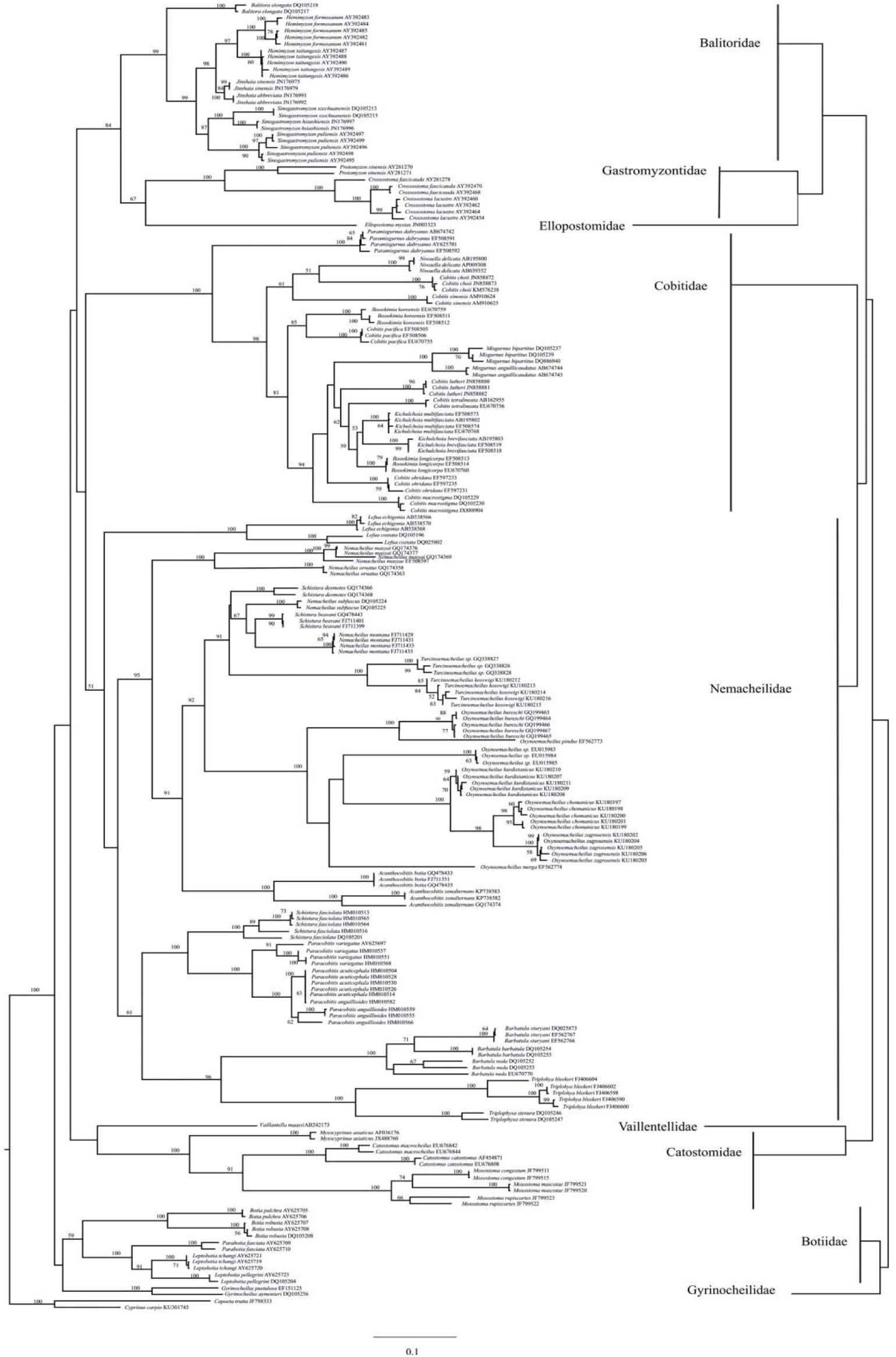
Phylogenetic tree relationships among taxa of the Cobitoidea, Branch lengths are proportional to inferred character substitutions under the GTR+G+I model. Numbers on branches are ML bootstrap values; those below 50% are not shown. An overview of the relationship obtained for Cobitoidea superfamily shown (right).

The reconstructed phylogenetic trees revealed the Cobitoidea superfamily as a monophyletic group (Fig 5 and 6). At the species level, all individuals were clustered in their monophyletic groups except one specimen of *Paracobitis anguillioides* that was grouped within the cluster of *Paracobitis acuticephala*. At the genus level, paraphyletic clusters were found for *Schistura*, *Nemacheilus*, *Cobitis*, *Iksookimia* and *Kichulchoia* genera (Fig. 5, 6). At the family level, all studied families were clustered in their monophyletic clades. Gyrinocheilidae and Botiidae represent the most-basal clades of the Cobitoidea superfamily and formed the sister group to the clade formed by Catostomidae, Vaillentellidae, Nemacheilidae, Cobitidae, Ellopostomidae, Gastromyzontidae and Balitoridae families. These trees showed monophyletic relationship between Nemacheilidae and Cobitidae families. The phylogenetic relationships indicated Balitoridae and Nemacheilidae as two distinctive families which had mistakenly been classified as one family.

## Discussion

Identification of loach sensu stricto species has always been difficult and their classification not stable (Siebert, 1987; Sawada, 1982; Berg, 1940; Hora, 1932; Regan, 1911). This presented a challenge for ichthyologist to undertake research on these species. The main problem to consider in studying loach species is identification and classification of cryptic and sibling species. Most problems appeared in two modes: differentiation between specimens from the same species and sometimes high similarity between specimens from different species. Molecular tools in combination with morphological methods were introduced to resolve these problems (Purry et al., 2016; Kelehear et al., 2011). The present study aimed to obtain molecular data for Choman loach species recently introduced based on morphological methods.

Results of univariate and multivariate analyses of truss network data greatly distinguished the four Choman loach species based on their truss distances. Univariate analysis showed that *O. zagrosensis* is more rotund than other Choman loach species. This specie is distinguished from other Choman loach species by having longer truss distances in the head, the trunk and the caudal peduncle parts of body. Multivariate analysis using DFAs well differentiated the four loach species based on their truss network data. Based on this analysis, the most discriminant landmarks were the distances located between fins, in the caudal fin, in the head region, in the trunk and in the caudal peduncle. Canonical scatter plot of DFA and dendrogram based on the cluster analysis using the most discriminative landmarks showed that *O. chomanicus* and *O. zagrosensis* are more similar but *O. zagrosensis* can be more easily distinguished through having longer landmarks than their corresponding landmarks in *O. chomanicus*. These results correspond to those reported for these two species using morphometric analysis (Kamangar et al., 2014).

The molecular analysis based on Cytochrome *b* sequences better identified the new loach species from the Choman sub-basin of Kurdistan, Iran. The topology of dendrograms obtained from the Cytochrome *b* sequences of the three new loach species were in accordance with the dendrogram obtained from the cluster analysis based on truss network measurements. Molecular data as well as morphometric analysis revealed three distinct new described species. We were unable to find similar sequences from *Oxynoemacheilus* in phylogenetic relationships by BA and ML trees which could be very close to the *Oxynoemacheilus* species from the Choman sub-basin. These results agree with the morphological classification of these species by Kamangar et al. (2014). *O. chomanicus* was originally described as a more similar species to *O. zagrosensis* based on morphological features (Kamangar et al., 2014). Two methods which have been broadly used for species identification based on DNA barcoding are genetic distance and monophyly based methods (Meyer et al., 2005; Zang, 2011; Zou et al., 2011). Two distinct monophyletic clades reconstructed based on Cytochrome *b* sequences of this new species within the Cobitoidea superfamily tree, did not show any overlapping between individuals from *O. zagrosensis* and *O. chomanicus*. Significant differences were also obtained in comparisons of K2P genetic distances between the mean intraspecific and the mean interspecies within the *Oxynoemacheilus* genus. The average K2P distances were 54.00-fold higher within the Nemacheilidae than that between individuals within the *Oxynoemacheilus* genus. These findings indicate the reliability of the Choman loach identification method based on Cytochrome *b* sequences and which were validated by Hebert et al. (2003) and Barrett & Hebert, (2005). The Cytochrome *b* sequences of *Turcinoemacheilus kosswigi* from the Choman river shows some similarity with *Turcinoemacheilus sp.* sequences as reported by Jamshidi et al. (2013) (Accessions; GQ 338826, GQ 338827 and GQ 338828). Jamshidi et al. (2013) later declared that these sequences belong to the *Turcinoemacheilus kosswigi.* Our findings did not confirm their declaration. Our reconstructed trees using BA and ML methods revealed that the *Turcinoemacheilus kosswigi* and *Turcinoemacheilus sp.* are two distinct species (Figs. 5 and 6). *Turcinoemacheilus hafezi* was also reported (Golzarianpour et al., 2013) more recently from the same area by Jamshidi et al. (2013) and it seems that these sequences are related to *T. hafezi*.

Cobitoidea is the most complicated fish group in regards their identification and phylogenetic relationship. It appears that to reach an acceptable phylogenic position for this superfamily will remain elusive. Our results clearly reveal that Cobitoidea is a monophyletic superfamily and comprises of nine lineages (Fig 5, 6). These classification results are in accordance with those reported by Šlechtová et al. (2007) in which Gyrinocheilidae, Botiidae, Vaillentellidae, Cobitidae, Catostomidae, Nemacheilidae and Balitoridae are valid family ranks. Moreover, our results illustrated that Ellopostomidae is a distinct family as previously described by Bohlen and Šlechtová (2009). Nemacheilidae and Balitoridae represent two distinct families as already described by Tang et al. (2006) and Šlechtová et al (2007). In our phylogenetic analysis, Nemacheilidae was clustered as a sister group with Cobitidae which was in agreement with those reported by Tang et al (2006) using mtDNA.

There are some disagreements regarding the Catostomidae phylogenetic relationship with Cobitoidea. Šlechtová et al. (2007) reported the Catostomidae as a distinct family within Cobitoidea based on molecular data using nuclear genes. However, some authors prefers to treat this group as a sister group with loaches sensu stricto (Tang et al., 2006; Chen et al., 2009) or even with other Cypriniformes (Mayden et al., 2009). The present results based on Cytochrome *b* demonstrate with strong statistical support that Catostomidae is a distinct monophyly family within Cobitoidea. These conflictions might be due to using different genes and their evolutionary histories as described by Mayden et al., 2009.

The Balitoridae cluster was divided into two distinct lineages. One cluster contains species that are currently classified as Balitoridae by some authors (Chen et al., 2009). Tang et al. (2006) treated this group as the Gastromyzoninae subfamily with Balitorinae in Balitoridae family. Kottelat (2012) classified the member of this cluster in the Gastromyzontidae family based on morphological analysis. Our results obtained strong statistical support for this lineage. When we included data from the Ellopostomidae family, the cluster of the Gastromyzontidae family showed a distinct lineage with high node support and as a sister group with the latter family. Thus, we consider this group as the Gastromyzontidae family as proposed morphologically by Kottelat (2012). Current analysis revealed that *Ellopostoma mystax* is a distinct lineage within the Cobitoidea superfamily and is more related to the Gastromyzontidae family than the Balitoridae family. Bohlen and Šlechtová (2009) introduced the Ellopostomidae family within the Cobitoidea superfamily as a valid family most closely related to the Nemacheilidae and Balitorodae families. However, they did not consider the family Gastromyzontidae in their analysis. Dissociating Gastromyzontidae from Balitoridae and including Ellopostmidae in one phylogenetic analysis could describe why former studies had difficulties in describing relationships of families in the Cobitoidea superfamily.

In conclusion, our molecular data shows the reliability of Choman loach species. The phylogentic analysis revealed nine distinct families within the Cobitoidea superfamily. However, we recommend more sample analysis of Barbuccidae and Serpenticobitidae to clarify complete molecular phylogeny of this superfamily.

## Author contributions

E. G., H. F., B. B. K. and M. A. N. designed the project, E. G. and B. B. K. conducted experiments; E. G., H. F. and B. B. K. analyzed the data; E. G. and B. B. K. wrote the manuscript; H. F. and B. B. K. edited the manuscript.

